# Involvement of orexin/hypocretin in the expression of social play behaviour in juvenile rats

**DOI:** 10.1101/809475

**Authors:** Christina J. Reppucci, Cassandra K. Gergely, Remco Bredewold, Alexa H. Veenema

## Abstract

Social play is a highly rewarding and motivated behaviour displayed by juveniles of many mammalian species. We hypothesized that the orexin/hypocretin (ORX) system is involved in the expression of juvenile social play behaviour because this system is interconnected with brain regions that comprise the social behaviour and mesocorticolimbic reward networks. We found that exposure to social play increased recruitment of ORX-A neurons in juvenile rats. Furthermore, central administration of ORX-A decreased social play duration, while central blockade of ORX-1 receptors differentially altered social play duration in juvenile rats with low versus high baseline levels of social play (increasing social play in low baseline social play individuals and decreasing social play in high baseline social play individuals). Together, our results provided the first evidence of a role for the ORX system in the modulation of juvenile social play behaviour.

## Introduction

Juvenile social play is a highly rewarding behaviour (Achterberg et al., 2016; Calcagnetti and Schechter, 1992; Humphreys and Einon, 1981) that can be modulated by motivational state (Ikemoto and Panksepp, 1992; Panksepp and Beatty, 1980), and is displayed by both sexes in many mammalian species (Bekoff and Byers, 1998; Pellis and Iwaniuk, 2000). This evolutionary conservation is likely due to the importance of social play in the development of social, emotional, and cognitive skills, which is essential for the expression of appropriate social interactions throughout life (Nijhof et al., 2018; Siviy, 2016; Spinka, Newberry, & Bekoff, 2001; van den Berg et al., 1999; Vanderschuren, Niesink, & VanRee, 1997). Understanding the neurobiological underpinnings of social play may have important implications for neurodevelopmental disorders like autism spectrum disorders (ASD), where patients often exhibit reduced motivation to engage in social play (Chevallier, Kohls, Troiani, Brodkin, & Schultz, 2012; Jordan, 2003).

We hypothesized that the orexin/hypocretin (ORX) system may be involved in the expression of social play behaviour, because this system is interconnected with brain regions that comprise the social behaviour and mesocorticolimbic reward networks (Newman, 1999; O’Connell and Hofmann, 2011), including regions previously implicated in the regulation of social play (e.g., Peyron et al., 1998; Schmitt et al., 2012; Vanderschuren, Achterberg, & Trezza, 2016). Although the ORX system has traditionally been associated with regulating the sleep-wake cycle and with stimulating feeding behaviour, emerging evidence has demonstrated its involvement in diverse motivated and reward-driven behaviours, especially in cases of high motivational relevance (for review see: Mahler, Moorman, Smith, James, & Aston-Jones, 2014). These diverse functions of the ORX system have been conceptualized to reflect its role in coordinating current motivational state with adaptive physiological and behavioural responses (for reviews see: Saper, 2006; Tsujino and Sakurai, 2009; Willie et al., 2003). However, whether the ORX system is involved in rewarding social behaviours, especially in juveniles, is largely unknown.

The ORX system is comprised of two neuropeptides (ORX-A, ORX-B) produced from the same prepro-ORX gene, and two receptors (ORX1R, ORX2R; de Lecea et al., 1998; Sakurai et al., 1998). Central synthesis of ORX is restricted to neurons in the lateral hypothalamic area (LHA) and adjacent dorsomedial and posterior hypothalamic nuclei (de Lecea, et al., 1998; Hahn, 2010; Sakurai, et al., 1998; Swanson, Sanchez-Watts, & Watts, 2005), but ORX projections are widespread throughout the brain (Peyron, et al., 1998; Schmitt, et al., 2012). Accordingly, ORX receptors are located in diverse brain regions (Marcus et al., 2001; Trivedi, Yu, MacNeil, Van der Ploeg, & Guan, 1998), and while ORX1R is preferentially activated by ORX-A, ORX2R has equal affinity for both neuropeptides (Sakurai, et al., 1998). ORX-A and ORX1Rs have been implicated in social behaviours in adult rodents (e.g., social interaction, maternal behaviors, male sexual behaviors; see: Abbas et al., 2015; Bai et al., 2009; D’Anna and Gammie, 2006; Di Sebastiano, Wilson-Perez, Lehman, & Coolen, 2011; Di Sebastiano, Yong-Yow, Wagner, Lehman, & Coolen, 2010; Muschamp, Dominguez, Sato, Shen, & Hull, 2007), and thus were selected as the focus of our current investigation.

We assessed the role of the ORX system in regulating social play in both male and female juvenile rats, because previous studies demonstrated sex-specific regulation of social play behaviour by other neuropeptide systems (Bredewold et al., 2018; Bredewold, Schiavo, van der Hart, Verreij, & Veenema, 2015; Bredewold, Smith, Dumais, & Veenema, 2014; Paul et al., 2014; Reppucci, Gergely, & Veenema, 2018; Veenema, Bredewold, & de Vries, 2013). We first examined whether juvenile rats exposed to social play had greater recruitment of ORX-A neurons than those not exposed to social play. We then examined whether central manipulations of ORX signalling altered the expression of social play behaviour. Because ORX signalling is often associated with promoting the expression of motivated or reward-driven behaviours (for review see: Mahler et al., 2014), we predicted enhanced recruitment of ORX-A neurons in response to social play exposure, and that central administration of ORX-A would increase the expression of social play while central blockade of ORX1Rs would decrease the expression of social play.

## Methods

### Subjects

Experimentally naïve juvenile male and female Wistar rats (Charles River Laboratories) were housed in single sex groups of two to four in standard rat cages (48 x 27 x 20 cm) and maintained under standard laboratory conditions (12 hr light/dark cycle, lights off at 14:00 h, food and water available *ad libitum*). All housing and testing was in accordance with the National Institute of Health *Guidelines for Care and Use of Laboratory Animals* and the Boston College and Michigan State University Institutional Animal Care and Use Committees.

### Social play testing and analysis

All experimental rats were individually housed in clean cages three days prior to the start of social play testing and were maintained individually housed for the remainder of each experiment (stimulus rats remained group housed). During social play testing, home cages were removed from the cage rack and placed on the floor of the housing room, wire cage lids were removed and replaced with a Plexiglas lid, and a tripod and video camera were set up above each cage. Tests lasted 10 min, during which time subjects in Social Play groups were exposed to an age- and sex-matched unfamiliar stimulus rat; subjects undergoing multiple social play tests (Experiments 2 and 3) received a different unfamiliar stimulus rat during each test. At the end of each test, the stimulus rat was removed, wire lids were replaced, and cages returned to the cage rack. Testing procedures were exactly as described above for subjects in No Social Play groups, except that they were not given access to a stimulus rat once their cage was placed on the floor. Stimulus rats were striped with a permanent marker 30-60 min prior to testing in order to distinguish them from the experimental rats during later video analysis. Food and water were not available during the 10 min testing sessions but were immediately returned when each session was complete. All testing took place during the first hour of the dark phase, testing order was counterbalanced across sex and testing conditions, and sessions were videotaped for future behavioural analyses. Subjects were 31-33 days old during testing (Figure 1).

**Figure 1.**
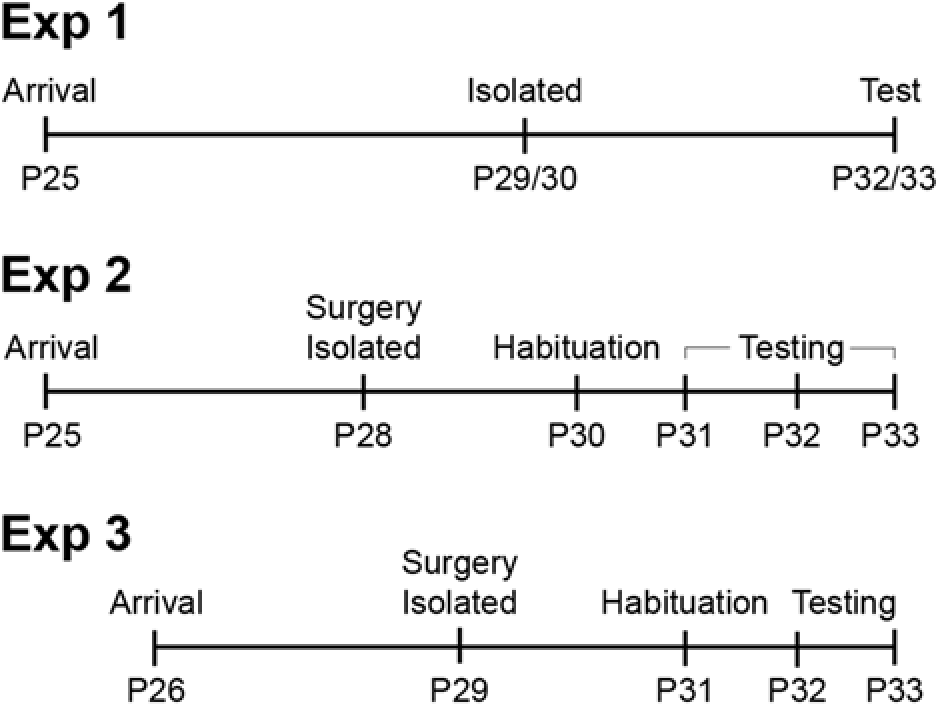
Experimental timelines. For all experiments, social play testing was conducted when rats were 31-33 days old. P = Postnatal day (age in days).

Behaviour was analysed using JWatcher (http://www.jwatcher.ucla.edu/) or Solomon Coder software (https://solomon.andraspeter.com/), by a trained observer unaware of the sex and testing condition of each subject. Videos were analysed for the percent of time that experimental subjects were engaged in social play (playful interactions with the stimulus rat), allogrooming (grooming the stimulus rat), and social investigation (e.g., sniffing the anogenital region of stimulus rat). The frequency of stereotypical social play elements, specifically, nape attacks (experimental rat attacks or makes nose contact with nape of stimulus rat), pins (experimental rat holds the stimulus rat in supine position), and supine positions (experimental rat is pinned on its back by the stimulus rat) was also analysed (see Supplemental Materials). Videos for subjects in the No Social Play groups were watched to confirm the presence of typical behaviours (such as cage exploration and self-grooming) and the absence of abnormal behaviours.

### Experiment 1: Activation of ORX-A neurons in response to social play exposure

After exposure to the 10-min social play test (n = 5 no social play condition, n = 8 social play condition; split evenly between the sexes), rats were left undisturbed in the housing room until sacrifice 80 min later at which time rats were deeply anesthetized with isoflurane, then transcardially perfused with 0.9% saline followed by 4% paraformaldehyde in 0.1 M borate buffer (pH 9.5; Figure 1). Brains were extracted and post-fixed 24 h in the perfusion solution, followed by 48 h in 30% sucrose, then rapidly frozen in 2-methylbutane cooled in dry ice, and stored at −45°C. Brains were sliced into 30μm coronal sections using a cryostat (Leica CM3050 S) and collected into three series which were stored in a cryoprotectant solution at −20°C until immunohistochemical processing. All histological procedures were completed in semi-darkness and out of direct light to preserve fluorescence.

One series of tissue was removed from cryoprotectant storage solution, thoroughly rinsed in tris-buffered saline solution (TBS; 50 mM, pH 7.6), and then processed sequentially, first for Fos (a commonly used marker of neuronal activation; Chaudhuri, 1997; Morgan and Curran, 1991) and then for ORX-A. To process for Fos, tissue was incubated for 24 h at 4°C in a blocking solution [TBS with 0.3% Triton X-100 and 2% normal donkey serum (017-000-121, Jackson ImmunoResearch)] with the primary antibody anti-cFos raised in rabbit (1:1K; SC-52; Santa Cruz Biotechnology Inc). After rinses in TBS, tissue was incubated for one hour in the blocking solution containing the secondary antibody Alexa 594 anti-rabbit raised in donkey (1:500; 711-585-152, Jackson ImmunoResearch). Following rinses in TBS, tissue was then processed for ORX-A by incubation for 24 h at 4°C in the blocking solution containing anti-orexin-A raised in goat (1:2K; sc-8070, Santa Cruz Biotechnology Inc). Tissue was then rinsed in TBS and incubated for 1 h in the blocking solution containing the secondary antibody Alexa 488 anti-goat raised in donkey (1:500; 705-545-147, Jackson ImmunoResearch). Following rinses in TBS, tissue was mounted onto slides, air-dried, coverslipped with Vectashield HardSet Mounting Medium with DAPI (4′,6-diamidino-2-phenylindole; nuclear counterstain; H-1500, Vector Labs), and stored at 4°C.

Images were acquired with a 20X objective on a Zeiss AxioImager fluorescence microscope with ApoTome attachment (Carl Zeiss Microscopy GmbH, Orca-R2 camera (Hamamatsu), and Zen software (Carl Zeiss Microscopy GmbH). Two bilateral sets of images were obtained per rat (Figure 2B) at three sampling locations: medial, perifornical, and lateral. Each image represented a sampling area that measured 433.45 µm x 330.25 µm. The number of double-labelled neurons [ORX-immunoreactive (-ir) + Fos-ir], the total number of ORX-ir neurons, and the total number of Fos-ir nuclei were counted as previously described (Petrovich, Hobin, & Reppucci, 2012; Reppucci, et al., 2018) and then averaged for each of the three sampling locations. The percent of ORX neurons that were co-labelled with Fos was quantified as: (# double-labelled neurons/total number of ORX-ir neurons)*100.

**Figure 2.**
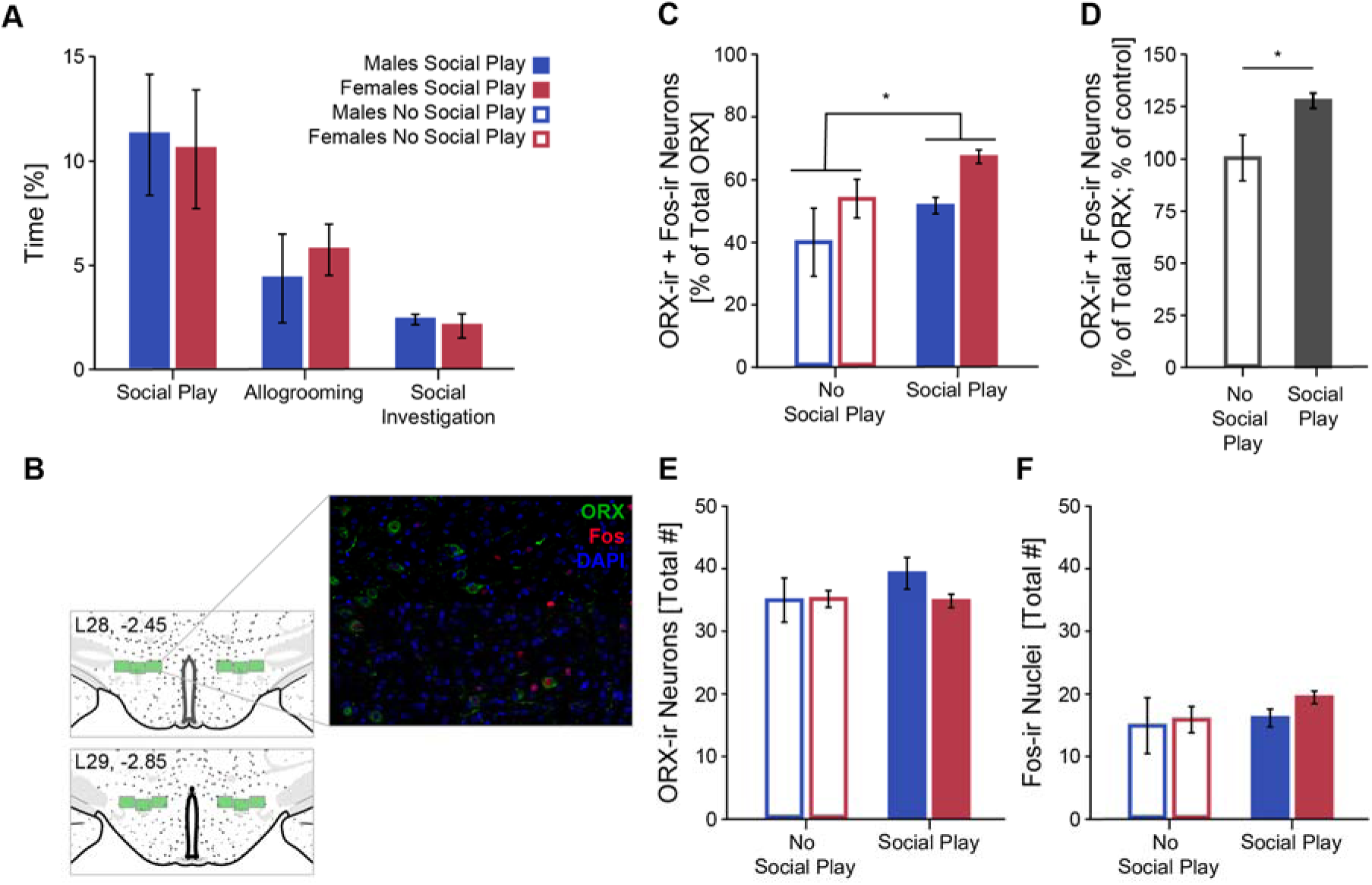
Exposure to social play increased recruitment of ORX-A neurons in juvenile rats. The percent of time juvenile rats spent engaged in social behaviours was similar between males and females (**A**). Fos induction within ORX-A neurons was quantified across the LHA (**B**; numbers refer to atlas figure (Swanson, 2018), distance from bregma in mm), and was higher in subjects exposed to social play than subjects in the no social play control condition (**C, D**). The total number of ORX-ir neurons (**E**) and Fos-ir nuclei (**F**) was similar across all groups. \**p* < 0.05, main effect of social play condition.

### Experiment 2: Effect of central administration of ORX-A on social play expression

While maintained under isoflurane anaesthesia (2-4%, as needed), subjects were implanted with a guide cannula (4mm, 21G; C312G/SPC, Plastics One) targeting the lateral ventricle (AP: −1.0, ML: - 1.6, DV: 2.0). Guide cannula were secured with three stainless steel screws (0-80 x 3/32, Plastics One) and dental cement (REFs 1404 & 1230, Lang Dental Mfg. Co.), and were closed with a dummy cannula (C312DC/1/SPC, Plastics One). Subjects received once daily subcutaneous injections of Rimadyl (10mg/kg; Henry Schein) on the day of surgery and the subsequent two recovery days before starting testing.

After recovery, rats were habituated to the social play testing procedures by receiving mock infusions (dummy cannula were removed/replaced and the motorized syringe pump was run) followed by social play testing 20 min later. Social play behaviour during habituation was not used in any analyses. On the subsequent 3 days, rats received, in a counterbalanced order, intracerebroventricular (ICV) infusions (3 µL) of vehicle (0.9% saline) or one of two doses of ORX-A (Sigma-Aldrich, O6012-.5MG; 0.1 or 1.0 nmol) into the lateral ventricle 20 min prior to each social play test (Figure 1). For infusions, dummy cannula were removed and an injector was inserted. Injectors were connected by tubing (PE50; C232CT, Plastics One) to 25 µL syringes (Hamilton Company, 7643-01) with 22G needles (Hamilton Company, 7748-08) that were mounted in a dual syringe pump (GenieTouch, Kent Scientific, Torrington, CT). Infusions were done over the course of 1 min, followed by 30 sec for diffusion from the injector tip. At the end of the experiment rats were euthanized using CO2 inhalation, blue food colouring diluted 50% in deionized water was injected into the guide cannula, brains were extracted, and then coronolly cut with a razor blade to verify intraventricular staining. Final groups sizes were n = 6/sex.

### Experiment 3: Effect of central blockade of ORX1Rs on social play expression

All procedures were as described in Experiment 2, except that subjects received counterbalanced ICV infusions (3 uL) of either vehicle [2% DMSO (Sigma-Aldrich, D8418-50ML) + 10% 2-hydroxypropyl-β-cyclodextrin (Sigma-Aldrich, H5784-10ML) in 0.9% saline] or the ORX1R antagonist SB-334867 (50 nmol; Tocris, 1960) into the lateral ventricle 20 min prior to social play testing (Figure 1). Final group sizes were n = 8 males and n = 10 females.

### Statistical analysis

For Experiment 1, independent sample t-tests were used to analyse the effect of sex on social behaviours, mixed-model [sampling region (within-subjects factor) x sex (between-subjects factor) x social play condition (between-subjects factor)] analysis of variances (ANOVAs) were used to assess the effects of sex and social play exposure on the activation of ORX-A neurons and a Pearson correlation (collapsed across sampling region and sex to increase power) was used to determine whether activation of ORX-A neurons (as measured by percent of the same-sex control group) correlated with the percent of time subjects engaged in social play. For Experiments 2 and 3, mixed-model [sex (between-subjects factor) x drug (within-subjects factor)] ANOVAs were used to assess the effects sex and ORX-A or SB-334867 on social behaviours. To assess whether differences in baseline levels of social play affected responsiveness to central manipulations of ORX signalling, data from Experiments 2 and 3 were collapsed across the sexes (because no sex differences were observed) and then divided by a mean split into “low social play” and “high social play” groups (based on the level of social play under vehicle conditions; Experiment 2: low social play n = 5, high social play n =7; Experiment 3: low social play n = 8, high social play n = 10). The effects of baseline social play levels and drug on social behaviours were then analysed using mixed-model [baseline social play level (between-subjects factor) x drug (within-subjects factor)] ANOVAs. When significant interactions were found in the ANOVAs, Bonferroni *post hoc* pairwise comparisons were conducted to clarify the effects. For mixed-model ANOVAs, Mauchly’s Test of Sphericity was consulted, and the Greenhouse-Geisser correction used when needed. All data were analysed using IBM SPSS Statistics 24, and statistical significance was set at *p <* 0.05.

## Results

### Experiment 1: Social play exposure was associated with increased recruitment of ORX-A neurons

As expected, there were no sex differences in the percent of time juvenile rats engaged in social play (*t*(6) = 0.17, *p* = 0.87), allogrooming (*t*(6) = 0.56, *p* = 0.60), or social investigation (*t*(6) = 0.48, *p* = 0.65; Fig 2A) behaviours.

There was a main effect of sampling region for Fos induction within ORX-A neurons (medial > perifornical > lateral), total number of ORX neurons analysed (medial ≅ perifornical > lateral), and total number of Fos-positive nuclei observed (medial > perifornical > lateral), but sampling region did not interact with sex or social play condition (Table 1, Supplemental Figure 1A-C). Juvenile rats exposed to social play had significantly greater Fos induction within ORX-A neurons compared to juvenile rats in the no social play control condition (Table 1, Figure 2C). Additionally, females exhibited significantly greater Fos induction within ORX-A neurons than males, irrespective of social play condition (Table 1, Figure 2C). To further examine this sex difference, the data were collapsed across sampling regions and transformed as a percent of the same-sex control group [each value computed as: (% of activated ORX neurons/mean % of activated ORX neurons for the control group of the same sex)*100]. This transformation eliminated the sex difference in the activation of ORX-A neurons (*F*(1,13) = 0.051, *p* = 0.83), but preserved the main effect of social play condition (*F*(1,13) = 6.71, *p* = 0.029; Figure 2D). However, the percent of time subjects engaged in social play was not significantly correlated with Fos induction within ORX neurons (*r*(8) = 0.34, *p* = 0.41).

**Table 1.** ANOVA statistics for Experiment 1: Activation of ORX-A neurons in response to social play exposure. Significant effects are indicated in **bold**.

|  | ORX-ir + Fos-ir<br>[% of Total ORX-ir] | ORX-ir [Total #] | Fos-ir [Total #] |
| --- | --- | --- | --- |
| Sampling Region | $F_{(2,18)} = \mathbf{65.8}, p < \mathbf{0.001}$ | $F_{(2,18)} = \mathbf{56.9}, p < \mathbf{0.001}$ | $F_{(2,18)} = \mathbf{80.7}, p < \mathbf{0.001}$ |
| Sampling Region x Sex | $F_{(2,18)} = 2.04, p = 0.16$ | $F_{(2,18)} = 0.85, p = 0.45$ | $F_{(2,18)} = 1.34, p = 0.29$ |
| Sampling Region x<br>Social Play Condition | $F_{(2,18)} = 0.77, p = 0.48$ | $F_{(2,18)} = 0.34, p = 0.72$ | $F_{(2,18)} = 0.76, p = 0.45$ |
| Sampling Region x Sex x<br>Social Play Condition | $F_{(2,18)} = 0.41, p = 0.67$ | $F_{(2,18)} = 0.24, p = 0.79$ | $F_{(2,18)} = 0.46, p = 0.64$ |
| Sex | $F_{(1,9)} = \mathbf{10.9}, p < \mathbf{0.001}$ | $F_{(1,9)} = 0.97, p = 0.35$ | $F_{(1,9)} = 1.20, p = 0.30$ |
| Social Play Condition | $F_{(1,9)} = \mathbf{8.67}, p = \mathbf{0.016}$ | $F_{(1,9)} = 0.86, p = 0.38$ | $F_{(1,9)} = 1.51, p = 0.25$ |
| Sex x Social Play Condition | $F_{(1,9)} = 0.012, p = 0.92$ | $F_{(1,9)} = 1.13, p = 0.32$ | $F_{(1,9)} = 0.35, p = 0.57$ |

Group differences in the activation of ORX-A neurons could not be explained by differences in the number of ORX-A neurons that were analysed, because the total number of ORX-A neurons were similar across all groups (Table 1, Figure 2E). Further, these differences were specific to ORX-A neurons, because the total number of Fos-ir nuclei was similar across all groups (Table 1, Figure 2F).

### Experiment 2: Central administration of ORX-A decreased social play expression

Central administration of ORX-A significantly decreased the percent of time juvenile rats engaged in social play (Table 2, Figure 3A) and allogrooming (Table 2, Figure 3B) behaviours in both males and females; there was no main effect of interaction of sex on either behavioural measure (Table 2). The percent of time juvenile rats spent investigating the stimulus animal was low and similar across all conditions and in both sexes (Table 2, Figure 3C).

**Figure 3.**
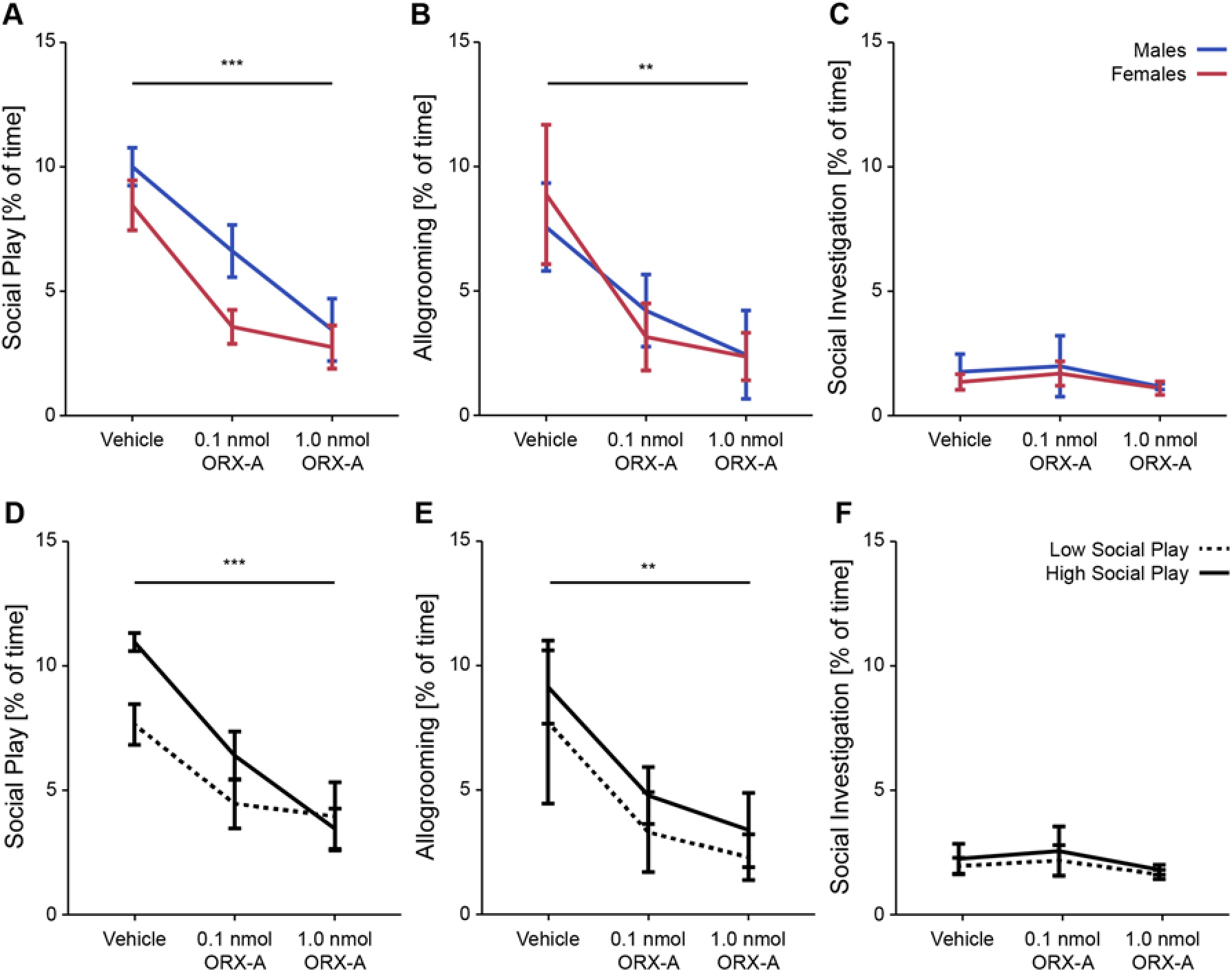
Central administration of ORX-A decreased social play expression in juvenile rats. Central administration of ORX-A decreased the expression of social play (**A**) and allogrooming (**B**), but did not affect social investigation (**C**). Central administration of ORX-A similarly affected subjects with low and high baseline levels of social play (**D-F**). *\*\*p* < 0.01, \*\*\**p* <0.001, main effect of drug.

**Table 2.**
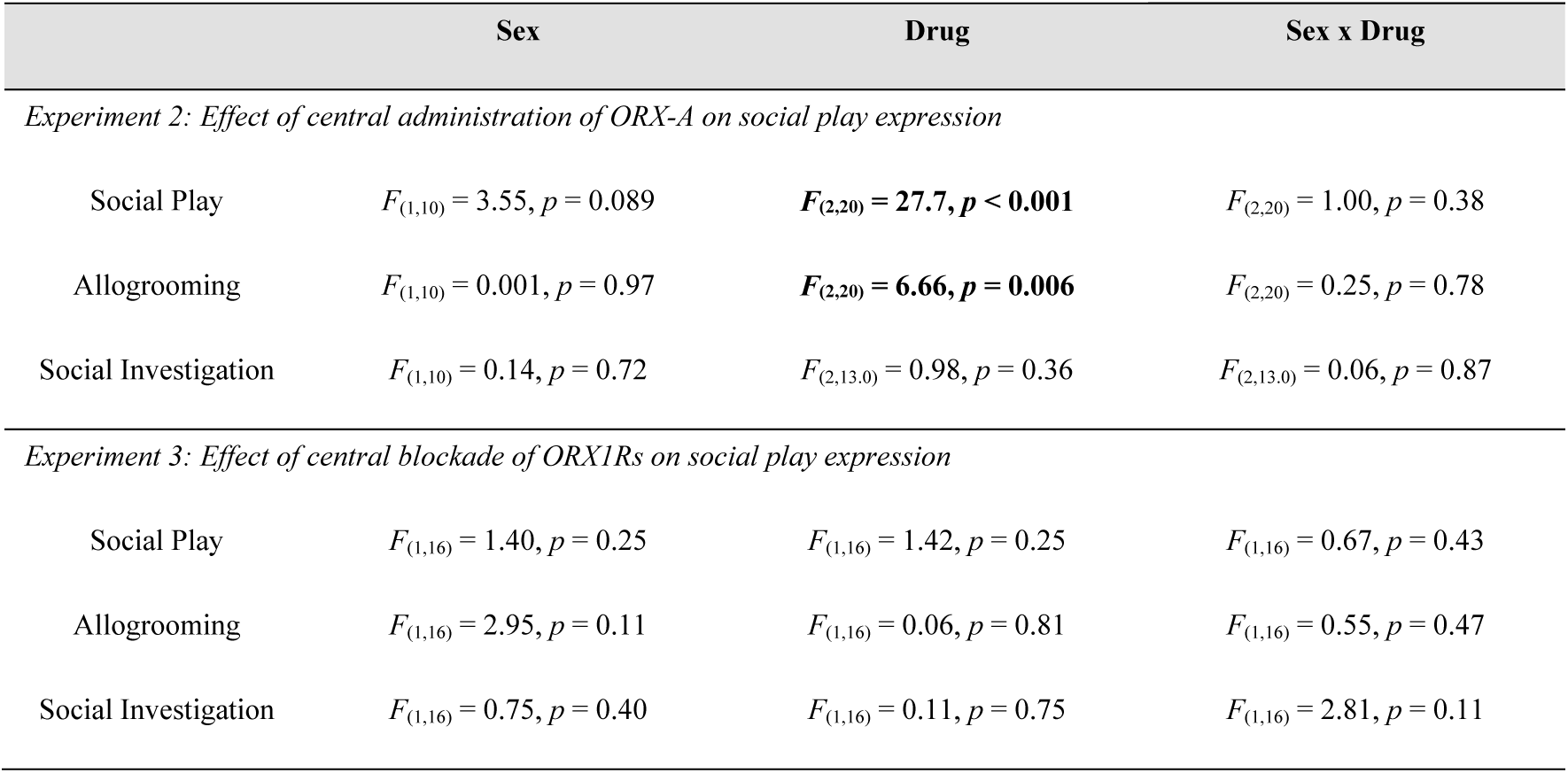
ANOVA statistics examining the effects of sex and central manipulations of ORX signalling on the expression of social behaviours in Experiments 2 and 3. Significant effects are indicated in **bold**.

The effect of ORX-A on social play expression was similar across all individuals (Figure 4A, Supplemental Figure 2A), and central administration of ORX-A similarly affected juvenile rats with low and high baseline levels of social play (Table 3, Figure 3D-E).

**Figure 4.**
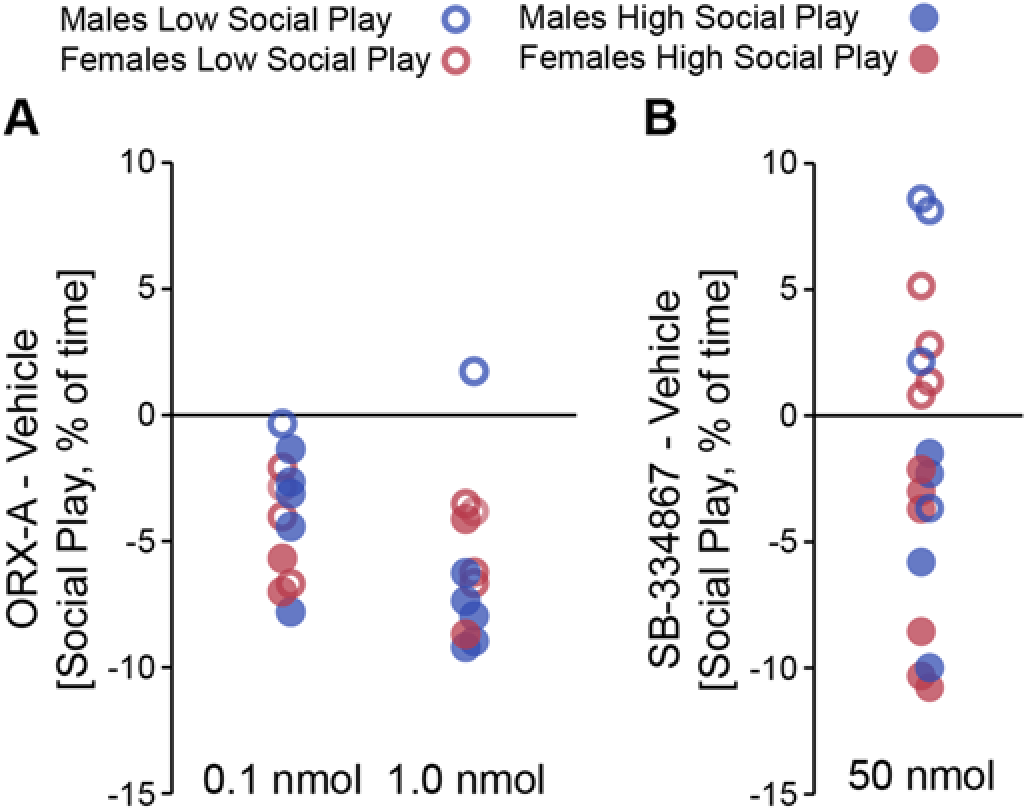
Individual changes in social play duration in response to central manipulations of ORX signalling. Graphical depiction of individual differences in the change in social play duration between vehicle and ORX-A in Experiment 2 (**A**) and between vehicle and ORX1R antagonist SB-334867 in Experiment 3 (**B**).

**Table 3.** ANOVA statistics examining how baseline differences in social play affected responsiveness to central manipulations of ORX signalling in Experiments 2 and 3. Significant effects are indicated in **bold**.

|  | Baseline Social Play Level | Drug | Baseline Social Play Level x<br>Drug |
| --- | --- | --- | --- |
| *Experiment 2: Effect of central administration of ORX-A on social play expression* |  |  |  |
| Social Play | $F_{(1,10)} = 2.81, p = 0.13$ | $F_{(2,20)} = \mathbf{29.5}, p < \mathbf{0.001}$ | $F_{(2,20)} = 3.35, p = 0.056$ |
| Allogrooming | $F_{(1,10)} = 0.77, p = 0.40$ | $F_{(2,20)} = \mathbf{6.26}, p = \mathbf{0.008}$ | $F_{(2,20)} = 0.008, p = 0.992$ |
| Social Investigation | $F_{(1,10)} = 0.19, p = 0.68$ | $F_{(2,20)} = 0.88, p = 0.43$ | $F_{(2,20)} = 0.018, p = 0.94$ |
| *Experiment 3: Effect of central blockade of ORX1Rs on social play expression* |  |  |  |
| Social Play | $F_{(1,16)} = 2.72, p = 0.12$ | $F_{(1,16)} = 2.08, p = 0.17$ | $F_{(1,16)} = \mathbf{23.70}, p < \mathbf{0.001}$ |
| Allogrooming | $F_{(1,16)} = 0.040, p = 0.85$ | $F_{(1,16)} = 0.054, p = 0.82$ | $F_{(1,16)} = 0.34, p = 0.57$ |
| Social Investigation | $F_{(1,16)} = 0.36, p = 0.56$ | $F_{(1,16)} = 0.065, p = 0.80$ | $F_{(1,16)} = 1.10, p = 0.31$ |

### Experiment 3: Central blockade of ORX1Rs differentially affected juvenile rats with low and high baseline levels of social play expression

Initial analyses indicated that central administration of the ORX1R antagonist SB-334867 did not alter the percent of time juvenile rats engaged in social play (Table 2, Figure 5A), allogrooming (Table 2, Figure 5B), or social investigation (Table 2, Figure 5C) in either sex; there was no main effect or interaction with sex on any behavioural measure (Table 2).

**Figure 5.**
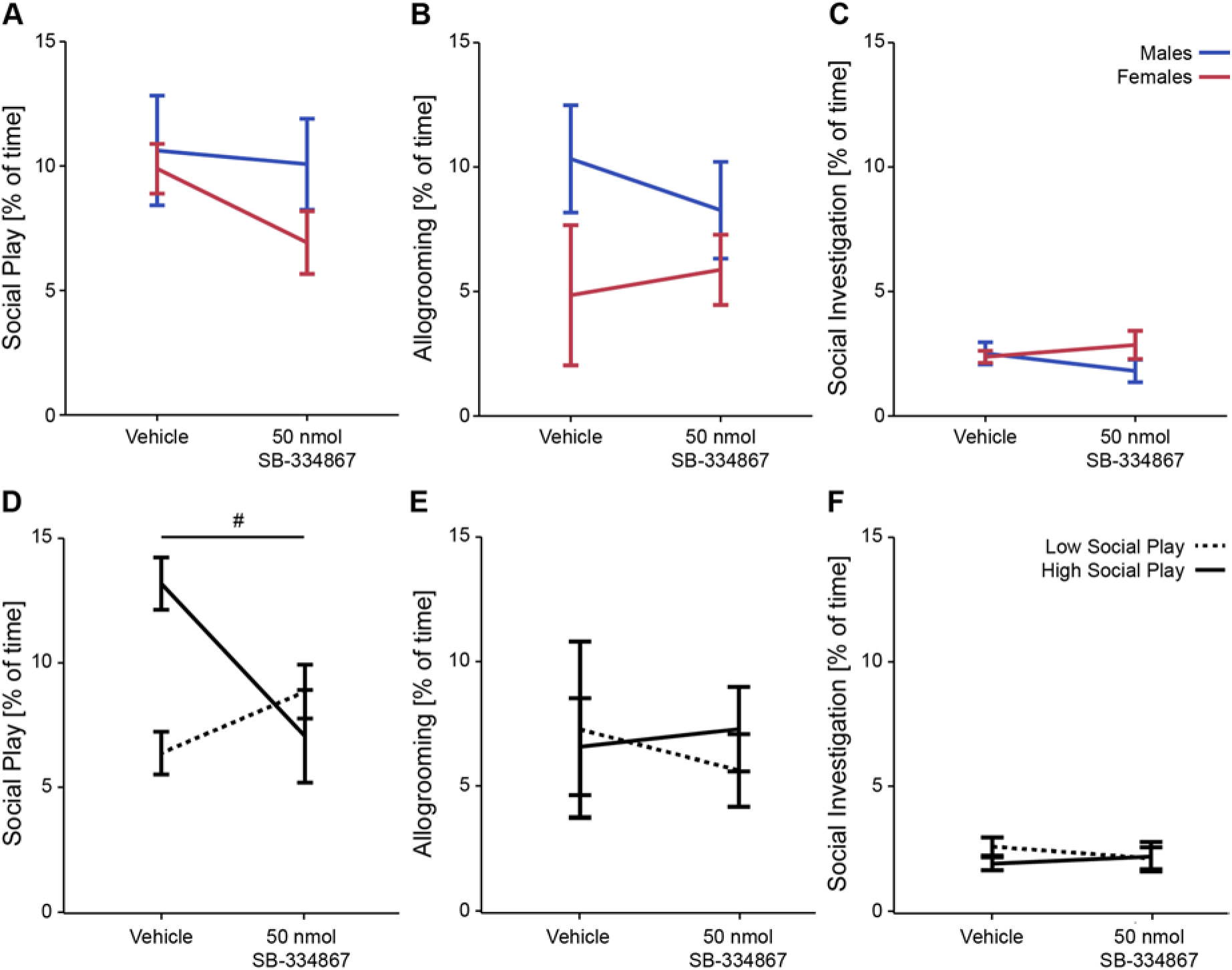
Central blockade of ORX1Rs differentially affected juvenile rats with low and high baseline levels of social play expression. Initial analyses indicated that central administration of the ORX1R antagonist SB-334867 did not alter the expression of social play in juvenile rats (**A**), however, follow-up analyses showed that SB-334867 differentially affected the expression of social play in subjects with low and high baseline levels of social play (**D**). Central administration of the ORX1R antagonist SB-334867 did not alter allogrooming (**B, E**) or social investigation (**C, F**) in any group. #*p* < 0.001, baseline social play level x drug interaction.

However, the effect of ORX1R blockade on social play expression varied across individuals (Figure 4B, Supplemental Figure 2B), and central administration of the ORX1R antagonist SB-334867 differentially affected juvenile rats with low and high baseline levels of social play as reflected by a significant baseline social play level (low vs. high levels of social play under vehicle) x drug interaction (Table 3, Figure 5D). Bonferroni *post hoc* paired comparisons showed that administration of SB-334867 significantly increased social play expression in subjects with low baseline levels of social play (*p* = 0.035) and significantly decreased social play expression in subjects with high baseline levels of social play (*p* < 0.001), such that both groups exhibited similar levels of social play when tested under SB-334867 (*p* = 0.38). The percent of time juvenile rats spent engaged in allogrooming and social investigation was similar across groups and conditions (Table 3, Figure 5E-F).

## Discussion

Here, we provided the first evidence for the involvement of the ORX system in the expression of social play behaviour in juvenile rats. In agreement with our predictions, exposure to social play increased the recruitment of ORX-A neurons. In contrast with our predictions, central administration of ORX-A decreased the expression of social play behaviour. Lastly, central blockade of ORX1Rs differentially altered social play behaviour in juvenile rats with low and high baseline levels of social play: increasing social play in low baseline social play subjects and decreasing social play in high baseline social play subjects. Together, these results suggest that the ORX system modulates the expression of social play behaviour in juvenile rats.

Increased Fos induction within ORX-A neurons following exposure to social play is consistent with studies investigating the involvement of the ORX system in other rewarding or motivated behaviours. Indeed, prior work demonstrated increased Fos induction within ORX neurons following exposure to cues for food (Campbell et al., 2017; Choi, Davis, Fitzgerald, & Benoit, 2010; Harris, Wimmer, & Aston-Jones, 2005; Petrovich, et al., 2012) or drugs (Martin-Fardon, Cauvi, Kerr, & Weiss, 2018; Richardson and Aston-Jones, 2012), and in response to pharmacologically-stimulated feeding (Nishimura et al., 2014; Silva et al., 2017). These data are also consistent with prior reports of increased ORX-A levels and ORX-A neuronal activation in response to social behaviours across mammalian species. For example, ORX-A levels increased in the amygdala with positive emotion and social interaction in humans (Blouin et al., 2013) and in cerebrospinal fluid with social play in dogs (Wu, Nienhuis, Maidment, Lam, & Siegel, 2011). Fos induction within ORX-A neurons also increased with mating (Muschamp, et al., 2007) and with exposure to a mating-paired conditioned place preference chamber (Di Sebastiano, et al., 2011) in adult male rats. Together, these data suggest that enhanced recruitment of ORX-A neurons is associated with the expression of rewarding behaviours, including rewarding social interactions, and that this role may be conserved across species.

We observed relatively high baseline levels of Fos induction within ORX neurons in the no social play control condition, which could be due to the cage handling procedures associated with testing and/or the time of testing. Regarding the latter, the social play test was conducted during the first hour of the dark phase, which represents the active phase in nocturnal species like laboratory rats (Siegel, 1961; Stephan and Zucker, 1972). ORX is well-known for its role in arousal and in modulating the sleep-wake cycle (for reviews see: Azeez, Del Gallo, Cristino, & Bentivoglio, 2018; Berridge, Espana, & Vittoz, 2010; Boutrel, Cannella, & de Lecea, 2010), and prior research found that Fos induction within ORX-A neurons was greater during the dark phase compared to the light phase (Estabrooke et al., 2001). Further, Fos induction within ORX-A neurons and ORX-A levels in the cerebrospinal fluid was greater during periods of wakefulness compared to periods of rest (Furlong, Vianna, Liu, & Carrive, 2009). Thus, the high baseline levels of ORX-A neuron activation in the no social play condition in the current study likely represents neuronal activity due to increased arousal and wakefulness associated with the onset of the active phase.

We observed a baseline sex difference in the activation of ORX-A neurons, with greater Fos induction within ORX-A neurons in females compared to males in both the social play and no social play conditions, however sample sizes were low. We hypothesize that this baseline sex difference in ORX-A activation could represent differences between males and females in their level of responsiveness to light cycle changes or circadian rhythms tied to light cycle changes (for review see: Bailey and Silver, 2014). Notably, plasma corticosterone levels peak at the start of the active phase, and this peak is higher in females than in males (Bailey and Silver, 2014). ICV administration of ORX-A has been shown to increase plasma corticosterone levels in males (Jaszberenyi, Bujdoso, Pataki, & Telegdy, 2000) and females (Moreno, Perello, Gaillard, & Spinedi, 2005), however, to our knowledge, no study has directly compared this effect between the sexes. Thus, whether the observed baseline sex difference in ORX-A activation could be linked to sex differences in HPA-axis activity during the start of the active phase is a question for future research.

While prepro-ORX mRNA (Jöhren, Neidert, Kummer, & Dominiak, 2002) and ORX-A protein (Taheri, Mahmoodi, Opacka-Juffry, Ghatei, & Bloom, 1999) levels have been shown to be higher in adult female compared to adult male rats using microdissection procedures, we did not observe a sex difference in the number of ORX-A neurons. This discrepancy could be due to the differences in the age of our test subjects. However, prior evidence suggests that ORX system development is at adult levels by 20-30 days of age in rats (Iwasa, Matsuzaki, Munkhzaya, Tungalagsuvd, Kuwahara, Yasui, & Irahara, M., 2015; Sawai, Ueta, Nakazato, & Ozawa, 2010; Yamamoto et al., 2000). Alternatively, this discrepancy could represent differences in sampling procedures, i.e., ORX content versus ORX-A neuron number. In support, our findings align with another group observing no sex differences in the number of ORX-A neurons (Funabashi et al., 2009). Thus, the results from the microdissection procedures could represent greater ORX content per neuron rather than a greater number of ORX-positive neurons in females compared to males.

Which aspect of the social play experience in the current experiment led to Fos induction within ORX-A neurons is unknown. In future experiments, additional control groups could help isolate the contribution of novel stimulus investigation (e.g., novel object investigation group) and physical activity (e.g., running wheel group) to the Fos induction patterns observed in Experiment 1. Further, exposure to a stimulus female (irrespective of female receptivity or the expression of mating behaviour) increased Fos induction within ORX-A neurons in sexually naïve and sexually experienced adult male rats (Di Sebastiano, et al., 2010) suggesting the exposure to a social stimulus alone may be enough to induce Fos expression within ORX-A neurons. Thus, to address whether ORX signalling influences the expression of social play behaviour, we next conducted central manipulations of ORX signalling and assessed the effects of these manipulations on the expression of social play.

In contrast to our prediction, we observed decreased expression of social play behaviour following ICV administration of ORX-A, and this effect was irrespective of baseline levels of social play. This unexpected result was unlikely due to the selected doses, because similar doses of ORX-A administered ICV were shown to increase food intake (e.g., Moreno, et al., 2005; Parise et al., 2011) and increase operant responding for food (e.g., Choi, et al., 2010; Kay, Parise, Lilly, & Williams, 2014). Instead, this finding could reflect specific roles for ORX signalling in the regulation of social versus non-social behaviours, and/or a role for ORX in modulating the expression of multiple motivated behaviours. For example, central administration of ORX-A may have enhanced food-directed motivation at the expense of social-directed motivation, despite the absence of food in the home cage during testing. Future experiments where subjects are given the choice to engage in social interaction versus food consumption or social contact-seeking versus food-seeking behaviours, would provide valuable insights into the role of the ORX system in modulating the expression of social versus non-social behaviours.

The role of ORX signalling in modulating social play expression could also be brain region-specific. It is possible that ORX signalling in some brain regions facilitates the expression of social play, and that ORX signalling in other brain regions inhibits the expression of social play. Indeed, we have previously shown that central versus local manipulations of a different neuropeptide system, the vasopressin system, can have opposite effects on social play expression (Veenema, et al., 2013). Thus, the exogenous application of ORX-A in Experiment 2 may have been biased in activating regions that inhibit social play expression, or suppressing regions that facilitate social play expression. Alternatively, our results could suggest that the role of increased recruitment of ORX-A neurons in response to social play exposure in Experiment 1 was to constrain the expression of social play.

However, our results are consistent with decreased maternal behaviour in mice (D’Anna and Gammie, 2006) and decreased opposite sex preference in male rats (Bai, et al., 2009) following similar doses of ICV administered ORX-A, as well as with a prior study where optogenetic stimulation of ORX neurons decreased time spent investigating a caged conspecific by adult male rats in a three-chambered social interaction assay (Heydendael, Sengupta, Beck, & Bhatnagar, 2014). Intriguingly, our findings also complement recent work in humans suggesting that the ORX system may be dysregulated in ASD patients, a clinical population which often exhibits reduced motivation to engage in social play (Chevallier, et al., 2012; Jordan, 2003). Specifically, there have been case reports of higher plasma levels of ORX-A (Messina et al., 2018) and ORX-B (Kobylinska et al., 2019) in ASD patients compared to the general population. Together, these studies suggest that high levels of ORX signalling may act to suppress the expression of social behaviours, and that the ORX system could represent a potential biomarker or novel therapeutic target for ASD.

We predicted that central blockade of ORX1R signalling would reduce the expression of social play because ORX1R knockout mice exhibit reduced sociality (Abbas, et al., 2015) and systemic blockade of ORX1R signalling impaired copulatory behaviours in male rats (Muschamp, et al., 2007). Instead, our initial analysis suggested that ICV administration of the ORX1R-antagonist did not affect the expression of any of the social behaviours analysed (Figure 5A-C). This lack of an effect was unlikely due to the selected dose, as similar doses of the same antagonist were shown to reduce pharmacologically-stimulated food intake (e.g., Karasawa, Yakabi, Wang, & Tache, 2014; Zheng, Patterson, & Berthoud, 2007). However, at the individual subject level ICV application of the ORX1R-anatagonist administration did alter social play expression (see Figure 4 and Supplementary Figure 2), and follow-up analyses showed that the direction of this change depended on the baseline levels of social play that subjects exhibited. Specifically, central blockade of ORX1Rs decreased social play expression in juvenile rats with high baseline levels of social play, and increased social play expression in juvenile rats with low baseline levels of social play. This suggests that ORX signalling may be involved in regulating the level of social play expression.

The ORX system has previously been associated with natural variation in the expression of other behaviours. For example, in adult male rodents, individual differences in extinction of conditioned fear responses (Sharko, Fadel, Kaigler, & Wilson, 2017), motivation for cocaine (Pantazis, James, Bentzley, & Aston-Jones, 2019), intrinsic spontaneous physical activity (Perez-Leighton, Boland, Billington, & Kotz, 2013), and resiliency/susceptibility in social defeat paradigms (Chung, Kim, Kim, Kim, & Yoon, 2014; Grafe, Eacret, Dobkin, & Bhatnagar, 2018) have all been associated with differences in the activation or organization of the ORX system. Additionally, central administration of ORX-A reduced preference for a receptive female in sexually high-motivated but not sexually low-motivated adult male rats (Bai, et al., 2009). Thus, it is possible that the ORX system may also be associated with individual differences in the expression of social play behaviour.

Individual differences in social play expression have been documented both within (Pellis and Mckenna, 1992; Poole and Fish, 1976; Taylor, 1980) and between (Northcutt and Nwankwo, 2018; Siviy, Love, DeCicco, Giordano, & Seifert, 2003) rat strains, and one neural system that has been implicated in these differences is the dopamine system (Pellis and Mckenna, 1992; Siviy, Crawford, Akopian, & Walsh, 2011). The ORX system has reciprocal functional connections with the mesolimbic dopamine system (e.g., Korotkova, Sergeeva, Eriksson, Haas, & Brown, 2003; Linehan, Trask, Briggs, Rowe, & Hirasawa, 2015; Vittoz, Schmeichel, & Berridge, 2008), and it is anatomically positioned to directly influence its functioning. ORX neurons project to the ventral tegmental area (Fadel and Deutch, 2002; Peyron, et al., 1998), which has ORX1Rs (Ch’ng and Lawrence, 2015) through which ORX can act on dopaminergic and GABAergic neurons (Korotkova, et al., 2003). Thus, we hypothesize that ORX interactions with the dopamine system could underlie differences in the responsiveness to ORX1R blockade that we observed in the current experiment. In the future, it would be informative to determine if baseline differences in social play correspond to differences in the number or activation of ORX1R-expressing neurons in the ventral tegmental area or other regions within the mesolimbic dopamine system.

In conclusion, the present set of experiments provided the first evidence for the involvement of the ORX system in the expression of social play behaviour in juvenile rats. Enhanced recruitment of ORX-A neurons by social play exposure and a reduction in social play expression by central application of ORX-A suggests that high levels of ORX-A may act to suppress or constrain social play behaviour. Along with the finding that central blockade of ORX1Rs differentially affected subjects with low and high baseline levels of social play, our results indicate a modulatory role for ORX signalling in regulating the level of social play expression in juvenile rats. Future investigations should aim to determine how these central manipulations of ORX signalling affected Fos induction within ORX and ORX1R-expressing neurons in subjects exposed to social play, where in the brain ORX is acting, and what other neural systems (e.g., dopamine system) it may be interacting with to influence the expression of social play behaviour.

## Biographical note

**Christina J. Reppucci:** Christina J. Reppucci is a Postdoctoral Researcher in the Neurobiology of Social Behaviour Lab. Her research is focused on delineating the functional neural circuitry underlying motivated behaviours (e.g., social behaviour, feeding behaviour) using system neuroscience approaches, and assessing whether that circuitry is differentially recruited in males and females.

**Cassandra K. Gergely:** Cassanda K. Gergely was an Undergraduate Thesis Student in the Neurobiology of Social Behaviour Lab whose research focused on quantifying the activation of neuropeptide populations during juvenile social play behaviour.

**Remco Bredewold:** Remco Bredewold is a Senior Research Associate in the Neurobiology of Social Behaviour Lab. His research interests focus on the roles of vasopressin and oxytocin in the development of social behaviour. More specifically, he is studying juvenile social play and social recognition.

**Alexa H. Veenema:** Alexa H. Veenema is an Associate Professor of Psychology, and the Director of the Neurobiology of Social Behaviour Lab at Michigan State University. Her research focuses on the neural basis of social behaviour. Understanding the regulation of social behaviour is essential to gain insights in normal social functioning as well as in abnormal social functioning as observed in e.g. autism spectrum disorder, personality disorders, mood and anxiety disorders, and schizophrenia. Her ultimate goal is to understand and treat the causes of social behaviour deficits more effectively.

## Acknowledgements

We would like to thank Leigha A. Brown, Ashley Q. Chambers, Valerie M. Khaykin, Nara F. Nascimento, Suhana S. Posani, Grace S. Ro, Catherine L. Washington, and Dr. Katie E. Yoest for technical assistance, and Dr. Alexander W. Johnson for thoughtful discussion regarding our results.

## Supplemental Material

The frequency of stereotypical social play behaviours was analysed as previously described (Veenema, et al., 2013). Nape attacks and pins are considered to be measures of social play motivation (appetitive and consummatory, respectively), while supine positions considered an indication of social play receptivity (Ikemoto and Panksepp, 1992; Panksepp and Beatty, 1980; Vanderschuren, et al., 1997), and are less frequently expressed by experimental rats in our testing paradigm (Bredewold, et al., 2018; Bredewold, et al., 2015; Bredewold, et al., 2014; Reppucci, et al., 2018; Veenema, et al., 2013).

**Supplemental Table 1.** Frequency of stereotypical social play elements. Data displayed as mean ± SEM; see Supplemental Tables 2 and 3 for corresponding statistics.

|  |  | Nape Attacks | Pins | Supine Positions |
| --- | --- | --- | --- | --- |
| *Experiment 1: Activation of ORX-A neurons in response to social play exposure* |  |  |  |  |
| Males (n =4) | Social Play Condition | 35.0 ± 9.41 | 10.5 ± 4.37 | 2.75 ± 1.11 |
| Females (n =4) | Social Play Condition | 26.3 ± 3.84 | 9.38 ± 2.38 | 3.25 ± 2.93 |
| *Experiment 2: Effect of central administration of ORX-A on social play expression* |  |  |  |  |
| Males (n = 6) | Vehicle | 60.7 ± 4.51 | 31.7 ± 6.05 | 11.5 ± 2.88 |
| | 0.1 nmol ORX-A | 38.0 ± 8.28 | 24.7 ± 5.68 | 8.33 ± 2.22 |
| | 1.0 nmol ORX-A | 23.8 ± 8.55 | 9.67 ± 2.43 | 3.67 ± 1.58 |
| Females (n = 6) | Vehicle | 48.2 ± 7.04 | 18.8 ± 6.05 | 3.5 ± 1.93 |
| | 0.1 nmol ORX-A | 13.8 ± 4.16 | 11.5 ± 3.62 | 4.50 ± 1.38 |
| | 1.0 nmol ORX-A | 13.3 ± 6.15 | 8.50 ± 2.74 | 4.50 ± 0.96 |
| *Experiment 3: Effect of central blockade of ORX1Rs on social play expression* |  |  |  |  |
| Males (n = 8) | Vehicle | 66.4 ± 12.86 | 26.3 ± 8.54 | 1.88 ± 0.52 |
| | SB-334867 | 69.5 ± 13.35 | 27.5 ± 7.69 | 3.38 ± 1.03 |
| Females (n = 10) | Vehicle | 61.2 ± 6.21 | 21.7 ± 4.84 | 8.60 ± 1.56 |
| | SB-334867 | 45.5 ± 8.58 | 18.6 ± 4.73 | 6.10 ± 1.57 |

**Supplemental Table 2.** Supplemental t-Test statistics for Experiment 1: Activation of ORX-A neurons in response to social play exposure.

|  |  |
| --- | --- |
| Nape Attacks | $t_{(6)} = 0.86, p = 0.42$ |
| Pins | $t_{(6)} = 0.45, p = 0.67$ |
| Supine Positions | $t_{(6)} = 0.16, p = 0.88$ |

**Supplemental Table 3.** Supplemental ANOVA statistics examining the effects of sex and central manipulations of ORX signalling on the expression of stereotypical social play elements in Experiments 2 and 3. Significant effects are indicated in **bold**. *Bonferroni *post hoc* paired comparisons revealed that males exhibited more supine positions than females under vehicle (*p* = 0.044), and that only in males did administration of ORX-A decrease supine positions (0.1 nmol: *p* = 0.036, 1.0 nmol: *p* = 0.008); no other paired comparisons reached significance (*p* > 0.05, all).

|  | Sex | Drug | Sex x Drug |
| --- | --- | --- | --- |
| *Experiment 2: Effect of central administration of ORX-A on social play expression* |  |  |  |
| Nape Attacks | $F_{(1,10)} = 4.17, p = 0.068$ | $F_{(2,20)} = \mathbf{32.26}, p < \mathbf{0.001}$ | $F_{(2,20)} = 1.23, p = 0.32$ |
| Pins | $F_{(1,10)} = 4.01, p = 0.073$ | $F_{(2,20)} = \mathbf{7.43}, p = \mathbf{0.004}$ | $F_{(2,20)} = 1.32, p = 0.29$ |
| Supine Positions | $F_{(1,10)} = 3.39, p = 0.095$ | $F_{(2,20)} = 2.35, p = 0.12$ | $F_{(2,20)} = \mathbf{3.76}, p = \mathbf{0.041}^*$ |
| *Experiment 3: Effect of central blockade of ORX1Rs on social play expression* |  |  |  |
| Nape Attacks | $F_{(1,16)} = 1.90, p = 0.19$ | $F_{(1,16)} = 0.42, p = 0.53$ | $F_{(1,16)} = 0.94, p = 0.35$ |
| Pins | $F_{(1,16)} = 1.19, p = 0.29$ | $F_{(1,16)} = 0.02, p = 0.89$ | $F_{(1,16)} = 0.11, p = 0.74$ |
| Supine Positions | $F_{(1,16)} = \mathbf{12.4}, p = \mathbf{0.003}$ | $F_{(1,16)} = 0.14, p = 0.72$ | $F_{(1,16)} = 2.17, p = 0.16$ |

**Supplemental Table 4.** Frequency of stereotypical social play elements in subjects with low and high baseline levels of social play in Experiments 2 and 3. Data displayed as mean ± SEM; see Supplemental Table 5 for corresponding statistics.

|  |  | Nape Attacks | Pins | Supine Positions |
| --- | --- | --- | --- | --- |
| *Experiment 2: Effect of central administration of ORX-A on social play expression* |  |  |  |  |
| Low Social Play<br>(n = 5) | Vehicle | 46.8 ± 8.53 | 15.2 ± 4.60 | 4.60 ± 3.06 |
| | 0.1 nmol ORX-A | 17.4 ± 7.09 | 8.60 ± 2.20 | 6.40 ± 2.23 |
| | 1.0 nmol ORX-A | 16.0 ± 10.8 | 9.80 ± 3.89 | 4.60 ± 1.17 |
| High Social Play<br>(n = 7) | Vehicle | 59.9 ± 3.83 | 32.4 ± 5.78 | 9.57 ± 2.64 |
| | 0.1 nmol ORX-A | 32.0 ± 8.02 | 24.9 ± 4.92 | 6.43 ± 1.89 |
| | 1.0 nmol ORX-A | 20.4 ± 5.53 | 8.57 ± 1.54 | 3.71 ± 1.34 |
| *Experiment 3: Effect of central blockade of ORX1Rs on social play expression* |  |  |  |  |
| Low Social Play<br>(n = 8) | Vehicle | 40.6 ± 6.12 | 8.75 ± 2.42 | 4.13 ± 1.53 |
| | SB-334867 | 59.9 ± 9.38 | 25.9 ± 5.49 | 5.63 ± 1.74 |
| High Social Play<br>(n = 10) | Vehicle | 81.8 ± 5.98 | 35.7 ± 5.51 | 6.80 ± 1.74 |
| | SB-334867 | 53.2 ± 12.5 | 19.9 ± 6.53 | 4.30 ± 1.53 |

**Supplemental Table 5.** Supplemental ANOVA statistics examining how baseline differences in social play affected responsiveness to central manipulations of ORX signalling in Experiments 2 and 3. Significant effects are indicated in **bold**. *Bonferroni *post hoc* paired comparisons showed that administration of ORX-A did not alter the frequency of pinning behaviour in subjects with low baseline social play (*p* > 0.05, all), but in subjects with high baseline social play the 1.0 nmol dose significantly reduced the frequency of pins compared to vehicle (*p* = 0.003) or the 0.1 nmol dose (*p* = 0.007). #Bonferroni *post hoc* paired comparisons showed that administration of SB-334867 significantly increased the frequency of pins, but not nape attacks, in subjects with low baseline social play (pins: *p* = 0.040; nape attacks: *p* = 0.13), and significantly decreased the frequency of pins and nape attacks in subjects with high baseline social play (pins: *p* = 0.035; nape attacks: *p* = 0.016).

|  | Baseline Social Play Level | Drug | Baseline Social Play Level x<br>Drug |
| --- | --- | --- | --- |
| *Experiment 2: Effect of central administration of ORX-A on social play expression* |  |  |  |
| Nape Attacks | $F_{(1,10)} = 1.53, p = 0.25$ | $F_{(2,20)} = \mathbf{28.9}, p < \mathbf{0.001}$ | $F_{(2,20)} = 0.62, p = 0.55$ |
| Pins | $F_{(1,10)} = \mathbf{6.45}, p = \mathbf{0.029}$ | $F_{(2,20)} = \mathbf{7.05}, p = \mathbf{0.005}$ | $F_{(2,20)} = \mathbf{3.55}, p = \mathbf{0.048}^*$ |
| Supine Positions | $F_{(1,10)} = 0.36, p = 0.56$ | $F_{(2,20)} = 1.48, p = 0.25$ | $F_{(2,20)} = 1.56, p = 0.23$ |
| *Experiment 3: Effect of central blockade of ORX1Rs on social play expression* |  |  |  |
| Nape Attacks | $F_{(1,16)} = 2.79, p = 0.11$ | $F_{(1,16)} = 0.034, p = 0.57$ | $F_{(1,16)} = \mathbf{8.94}, p = \mathbf{0.009}^\#$ |
| Pins | $F_{(1,16)} = 3.23, p = 0.091$ | $F_{(1,16)} = 0.017, p = 0.90$ | $F_{(1,16)} = \mathbf{10.3}, p = \mathbf{0.005}^\#$ |
| Supine Positions | $F_{(1,16)} = 0.14, p = 0.71$ | $F_{(1,16)} = 0.14, p = 0.72$ | $F_{(1,16)} = 2.17, p = 0.16$ |

**Supplemental Figure 1.**
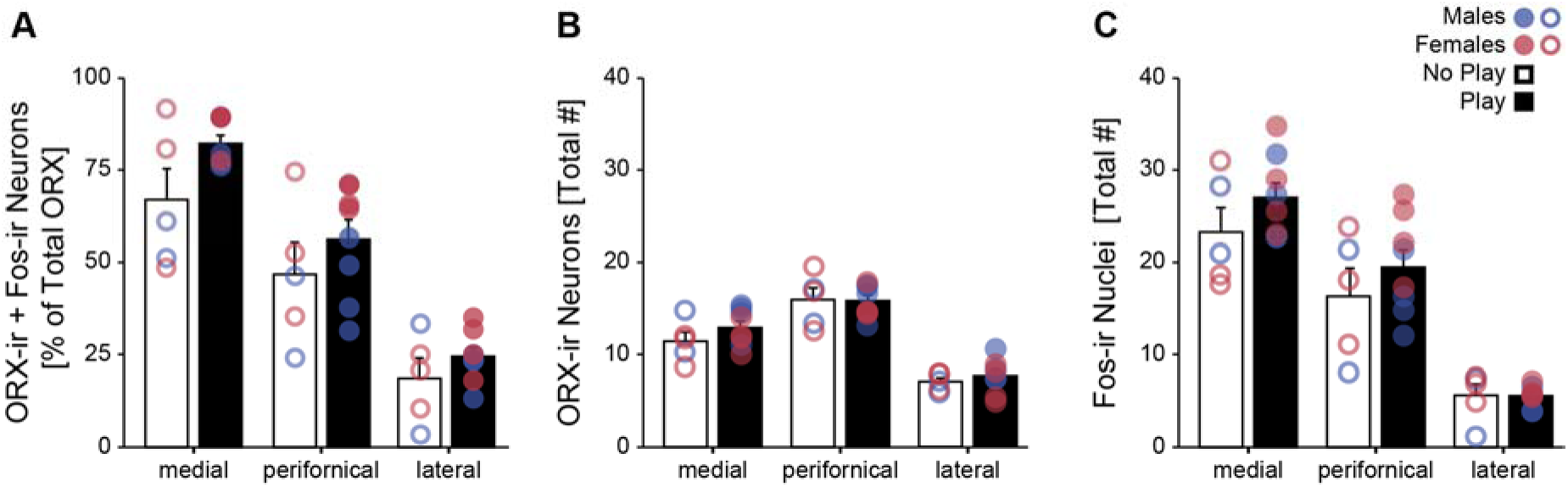
Topographical differences in ORX and Fos counts in Experiment 1. There was a main effect of sampling region for Fos induction within ORX-A neurons (**A**), the total number of ORX neurons analysed (**B**), and the total number of Fos-positive nuclei observed (**C**). However, sampling region did not interact with sex or social play condition, and thus, data were combined across sampling regions for presentation in Figure 2 (See Results).

**Supplemental Figure 2.**
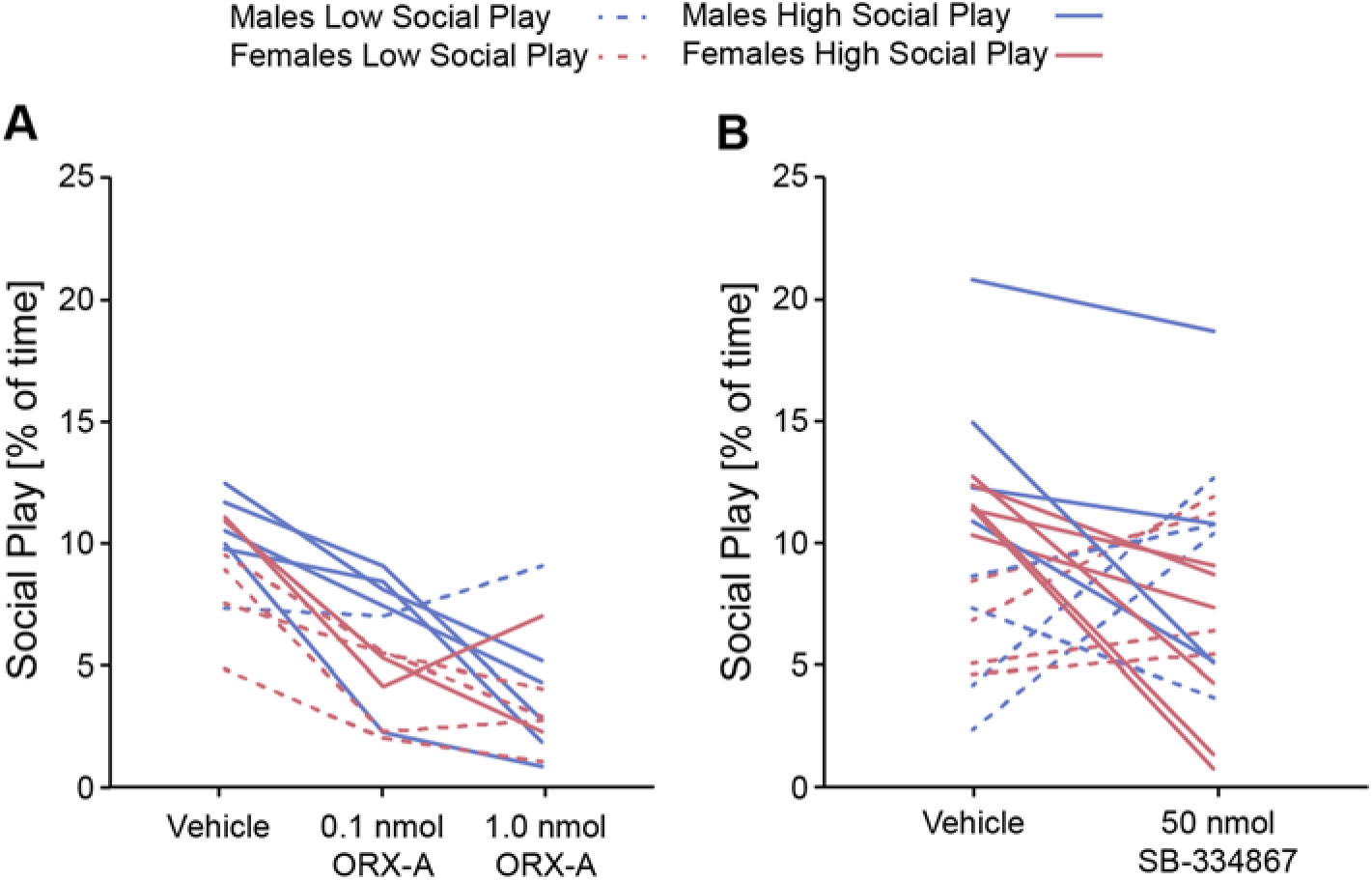
Individual changes in social play duration in response to central manipulations of ORX signalling. Individual data depicting a consistent decrease in social play duration in response to central administration of ORX-A in Experiment 2 (**A;** data from Figure 3A, D) and a bimodal change in social play duration in response to central administration of the ORX1R antagonist SB-334867 in Experiment 3 (**B;** data from Figure 5A, D).

## References

Abbas, M. G., Shoji, H., Soya, S., Hondo, M., Miyakawa, T., & Sakurai, T. (2015). Comprehensive behavioral analysis of male Ox1r (-/-) mice showed implication of orexin receptor-1 in mood, anxiety, and social behavior. Front Behav Neurosci, 9, p 324. doi:10.3389/fnbeh.2015.00324

Achterberg, E. J. M., van Kerkhof, L. W. M., Servadio, M., van Swieten, M. M. H., Houwing, D. J., Aalderink, M., … Vanderschuren, L. J. M. J. (2016). Contrasting roles of dopamine and noradrenaline in the motivational properties of social play behavior in rats. Neuropsychopharmacology, 41(3), pp. 858–868. doi:10.1038/npp.2015.212

Azeez, I. A., Del Gallo, F., Cristino, L., & Bentivoglio, M. (2018). Daily fluctuation of orexin neuron activity and wiring: The challenge of “chronoconnectivity”. Front Pharmacol, 9, p 1061. doi:10.3389/fphar.2018.01061

Bai, Y. J., Li, Y. H., Zheng, X. G., Han, J., Yang, X. Y., & Sui, N. (2009). Orexin A attenuates unconditioned sexual motivation in male rats. Pharmacol Biochem Behav, 91(4), pp. 581–589. doi:10.1016/j.pbb.2008.09.018

Bailey, M., & Silver, R. (2014). Sex differences in circadian timing systems: implications for disease. Front Neuroendocrinol, 35(1), pp. 111–139. doi:10.1016/j.yfrne.2013.11.003

Bekoff, M., & Byers, J. A. (1998). Animal Play: Evolutionary, Comparative, and Ecological Perspectives Cambridge: Cambridge University Press.

Berridge, C. W., Espana, R. A., & Vittoz, N. M. (2010). Hypocretin/orexin in arousal and stress. Brain Res, 1314, pp. 91–102. doi:10.1016/j.brainres.2009.09.019

Blouin, A. M., Fried, I., Wilson, C. L., Staba, R. J., Behnke, E. J., Lam, H. A., … Siegel, J. M. (2013). Human hypocretin and melanin-concentrating hormone levels are linked to emotion and social interaction. Nature Communications, 4, p 1547. doi:10.1038/ncomms2461

Boutrel, B., Cannella, N., & de Lecea, L. (2010). The role of hypocretin in driving arousal and goal-oriented behaviors. Brain Research, 1314, pp. 103–111.

Bredewold, R., Nascimento, N. F., Ro, G. S., Cieslewski, S. E., Reppucci, C. J., & Veenema, A. H. (2018). Involvement of dopamine, but not norepinephrine, in the sex-specific regulation of juvenile socially rewarding behavior by vasopressin. Neuropsychopharmacology, 43(10), p 2109. doi:10.1038/s41386-018-0100-2

Bredewold, R., Schiavo, J. K., van der Hart, M., Verreij, M., & Veenema, A. H. (2015). Dynamic changes in extracellular release of GABA and glutamate in the lateral septum during social play behavior in juvenile rats: Implications for sex-specific regulation of social play behavior. Neuroscience, 307, pp. 117–127. doi:10.1016/j.neuroscience.2015.08.052

Bredewold, R., Smith, C. J. W., Dumais, K. M., & Veenema, A. H. (2014). Sex-specific modulation of juvenile social play behavior by vasopressin and oxytocin depends on social context. Frontiers in Behavioral Neuroscience, 8(216). doi:10.3389/Fnbeh.2014.00216

Calcagnetti, D. J., & Schechter, M. D. (1992). Place conditioning reveals the rewarding aspect of social-interaction in juvenile rats. Physiology & Behavior, 51(4), pp. 667–672. doi:Doi 10.1016/0031-9384(92)90101-7

Campbell, E. J., Barker, D. J., Nasser, H. M., Kaganovsky, K., Dayas, C. V., & Marchant, N. J. (2017). Cue-induced food seeking after punishment is associated with increased Fos expression in the lateral hypothalamus and basolateral and medial amygdala. Behav Neurosci, 131(2), pp. 155–167. doi:10.1037/bne0000185

Ch’ng, S. S., & Lawrence, A. J. (2015). Distribution of the orexin-1 receptor (OX1R) in the mouse forebrain and rostral brainstem: A characterisation of OX1R-eGFP mice. J Chem Neuroanat, 66-67, pp. 1–9. doi:10.1016/j.jchemneu.2015.03.002

Chaudhuri, A. (1997). Neural activity mapping with inducible transcription factors. Neuroreport, 8(16), pp. iii–vii.

Chevallier, C., Kohls, G., Troiani, V., Brodkin, E. S., & Schultz, R. T. (2012). The social motivation theory of autism. Trends in Cognitive Sciences, 16(4), pp. 231–239. doi:10.1016/j.tics.2012.02.007

Choi, D. L., Davis, J. F., Fitzgerald, M. E., & Benoit, S. C. (2010). The role of orexin-A in food motivation, reward-based feeding behavior and food-induced neuronal activation in rats. Neuroscience, 167(1), pp. 11–20. doi:10.1016/j.neuroscience.2010.02.002

Chung, H. S., Kim, J. G., Kim, J. W., Kim, H. W., & Yoon, B. J. (2014). Orexin administration to mice that underwent chronic stress produces bimodal effects on emotion-related behaviors. Regulatory Peptides, 194, pp. 16–22. doi:10.1016/j.regpep.2014.11.003

D’Anna, K. L., & Gammie, S. C. (2006). Hypocretin-1 dose-dependently modulates maternal behaviour in mice. J Neuroendocrinol, 18(8), pp. 553–566. doi:10.1111/j.1365-2826.2006.01448.x

de Lecea, L., Kilduff, T. S., Peyron, C., Gao, X., Foye, P. E., Danielson, P. E., … Sutcliffe, J. G. (1998). The hypocretins: hypothalamus-specific peptides with neuroexcitatory activity. Proc Natl Acad Sci U S A, 95(1), pp. 322–327.

Di Sebastiano, A. R., Wilson-Perez, H. E., Lehman, M. N., & Coolen, L. M. (2011). Lesions of orexin neurons block conditioned place preference for sexual behavior in male rats. Horm Behav, 59(1), pp. 1–8. doi:10.1016/j.yhbeh.2010.09.006

Di Sebastiano, A. R., Yong-Yow, S., Wagner, L., Lehman, M. N., & Coolen, L. M. (2010). Orexin mediates initiation of sexual behavior in sexually naive male rats, but is not critical for sexual performance. Horm Behav, 58(3), pp. 397–404. doi:10.1016/j.yhbeh.2010.06.004

Estabrooke, I. V., McCarthy, M. T., Ko, E., Chou, T. C., Chemelli, R. M., Yanagisawa, M., … Scammell, T. E. (2001). Fos expression in orexin neurons varies with behavioral state. J Neurosci, 21, pp. 1656–1662.

Fadel, J., & Deutch, A. Y. (2002). Anatomical substrates of orexin-dopamine interactions: lateral hypothalamic projections to the ventral tegmental area. Neuroscience, 111(2), pp. 379–387. doi:10.1016/s0306-4522(02)00017-9

Funabashi, T., Hagiwara, H., Mogi, K., Mitsushima, D., Shinohara, K., & Kimura, F. (2009). Sex differences in the responses of orexin neurons in the lateral hypothalamic area and feeding behavior to fasting. Neuroscience letters, 463(1), pp. 31–34. doi: 10.1016/j.neulet.2009.07.035

Furlong, T. M., Vianna, D. M., Liu, L., & Carrive, P. (2009). Hypocretin/orexin contributes to the expression of some but not all forms of stress and arousal. Eur J Neurosci, 30(8), pp. 1603–1614. doi:10.1111/j.1460-9568.2009.06952.x

Grafe, L. A., Eacret, D., Dobkin, J., & Bhatnagar, S. (2018). Reduced orexin system function contributes to resilience to repeated social stress. Eneuro, 5(2). doi:10.1523/Eneuro.0273-17.2018

Hahn, J. D. (2010). Comparison of melanin-concentrating hormone and hypocretin/orexin peptide expression patterns in a current parceling scheme of the lateral hypothalamic zone. Neurosci Lett, 468(1), pp. 12–17. doi:10.1016/j.neulet.2009.10.047

Harris, G. C., Wimmer, M., & Aston-Jones, G. (2005). A role for lateral hypothalamic orexin neurons in reward seeking. Nature, 437(7058), pp. 556–559. doi:10.1038/nature04071

Heydendael, W., Sengupta, A., Beck, S., & Bhatnagar, S. (2014). Optogenetic examination identifies a context-specific role for orexins/hypocretins in anxiety-related behavior. Physiol Behav, 130, pp. 182–190. doi:10.1016/j.physbeh.2013.10.005

Humphreys, A. P., & Einon, D. F. (1981). Play as a reinforcer for maze-learning in juvenile rats. Animal Behaviour, 29(Feb), pp. 259–270. doi:10.1016/S0003-3472(81)80173-X

Ikemoto, S., & Panksepp, J. (1992). The effects of early social-isolation on the motivation for social play in juvenile rats. Developmental Psychobiology, 25(4), pp. 261–274. doi: 10.1002/dev.420250404

Iwasa, T., Matsuzaki, T., Munkhzaya, M., Tungalagsuvd, A., Kuwahara, A., Yasui, T., & Irahara, M. (2015). Developmental changes in the hypothalamic mRNA levels of prepro-orexin and orexin receptors and their sensitivity to fasting in male and female rats. International Journal of Developmental Neuroscience, 46, pp. 51–54. doi:10.1016/j.ijdevneu.2015.07.005

Jaszberenyi, M., Bujdoso, E., Pataki, I., & Telegdy, G. (2000). Effects of orexins on the hypothalamic-pituitary-adrenal system. J Neuroendocrinol, 12(12), pp. 1174–1178. doi:10.1046/j.1365-2826.2000.00572.x

Jöhren, O., Neidert, S. J., Kummer, M., & Dominiak, P. (2002). Sexually dimorphic expression of prepro-orexin mRNA in the rat hypothalamus. Peptides, 23(6), pp. 1177–1180. doi:10.1016/s0196-9781(02)00052-9

Jordan, R. (2003). Social play and autistic spectrum disorders - A perspective on theory, implications and educational approaches. Autism, 7(4), pp. 347–360. doi:10.1177/1362361303007004002

Karasawa, H., Yakabi, S., Wang, L., & Tache, Y. (2014). Orexin-1 receptor mediates the increased food and water intake induced by intracerebroventricular injection of the stable somatostatin pan-agonist, ODT8-SST in rats. Neurosci Lett, 576, pp. 88–92. doi:10.1016/j.neulet.2014.05.063

Kay, K., Parise, E. M., Lilly, N., & Williams, D. L. (2014). Hindbrain orexin 1 receptors influence palatable food intake, operant responding for food, and food-conditioned place preference in rats. Psychopharmacology (Berl), 231(2), pp. 419–427. doi:10.1007/s00213-013-3248-9

Kobylinska, L., Panaitescu, A. M., Gabreanu, G., Anghel, C. G., Mihailescu, I., Rad, F., & Zagrean, A. M. (2019). Plasmatic levels of neuropeptides, including oxytocin, in children with autism spectrum disorder, correlate with the disorder severity. Acta Endocrinol (Buchar), -5(1), pp. 16–24. doi:10.4183/aeb.2019.16

Korotkova, T. M., Sergeeva, O. A., Eriksson, K. S., Haas, H. L., & Brown, R. E. (2003). Excitation of ventral tegmental area dopaminergic and nondopaminergic neurons by orexins/hypocretins. J Neurosci, 23(1), pp. 7–11.

Linehan, V., Trask, R. B., Briggs, C., Rowe, T. M., & Hirasawa, M. (2015). Concentration-dependent activation of dopamine receptors differentially modulates GABA release onto orexin neurons. Eur J Neurosci, 42(3), pp. 1976–1983. doi:10.1111/ejn.12967

Mahler, S. V., Moorman, D. E., Smith, R. J., James, M. H., & Aston-Jones, G. (2014). Motivational activation: a unifying hypothesis of orexin/hypocretin function. Nat Neurosci, 17(10), pp. 1298–1303. doi:10.1038/nn.3810

Marcus, J. N., Aschkenasi, C. J., Lee, C. E., Chemelli, R. M., Saper, C. B., Yanagisawa, M., & Elmquist, J. K. (2001). Differential expression of orexin receptors 1 and 2 in the rat brain. J Comp Neurol, 435(1), pp. 6–25. doi:10.1002/cne.1190

Martin-Fardon, R., Cauvi, G., Kerr, T. M., & Weiss, F. (2018). Differential role of hypothalamic orexin/hypocretin neurons in reward seeking motivated by cocaine versus palatable food. Addict Biol, 23(1), pp. 6–15. doi:10.1111/adb.12441

Messina, A., Monda, V., Sessa, F., Valenzano, A., Salerno, M., Bitetti, I., … Carotenuto, M. (2018). Sympathetic, metabolic adaptations, and oxidative stress in autism spectrum disorders: How far from physiology? Front Physiol, 9, p 261. doi:10.3389/fphys.2018.00261

Moreno, G., Perello, M., Gaillard, R. C., & Spinedi, E. (2005). Orexin a stimulates hypothalamic-pituitary-adrenal (HPA) axis function, but not food intake, in the absence of full hypothalamic NPY-ergic activity. Endocrine, 26(2), pp. 99–106. doi:10.1385/ENDO:26:2:099

Morgan, J. I., & Curran, T. (1991). Stimulus-transcription coupling in the nervous system: involvement of the inducible proto-oncogenes fos and jun. Annu Rev Neurosci, 14, pp. 421–451. doi:10.1146/annurev.ne.14.030191.002225

Muschamp, J. W., Dominguez, J. M., Sato, S. M., Shen, R. Y., & Hull, E. M. (2007). A role for hypocretin (orexin) in male sexual behavior. J Neurosci, 27(11), pp. 2837–2845. doi:10.1523/JNEUROSCI.4121-06.2007

Newman, S. W. (1999). The medial extended amygdala in male reproductive behavior. A node in the mammalian social behavior network. Ann N Y Acad Sci, 877, pp. 242–257. doi:10.1111/j.1749-6632.1999.tb09271.x

Nijhof, S. L., Vinkers, C. H., van Geelen, S. M., Duijff, S. N., Achterberg, E. J. M., van der Net, J., … Lesscher, H. M. B. (2018). Healthy play, better coping: The importance of play for the development of children in health and disease. Neurosci Biobehav Rev, 95, pp. 421–429. doi:10.1016/j.neubiorev.2018.09.024

Nishimura, Y., Mabuchi, K., Taguchi, S., Ikeda, S., Aida, E., Negishi, H., & Takamata, A. (2014). Involvement of orexin-A neurons but not melanin-concentrating hormone neurons in the short-term regulation of food intake in rats. J Physiol Sci, 64(3), pp. 203–211. doi:10.1007/s12576-014-0312-0

Northcutt, K. V., & Nwankwo, V. C. (2018). Sex differences in juvenile play behavior differ among rat strains. Dev Psychobiol, 60(8), pp. 903–912. doi:10.1002/dev.21760

O’Connell, L. A., & Hofmann, H. A. (2011). The vertebrate mesolimbic reward system and social behavior network: a comparative synthesis. J Comp Neurol, 519(18), pp. 3599–3639. doi:10.1002/cne.22735

Panksepp, J., & Beatty, W. W. (1980). Social deprivation and play in rats. Behavioral and Neural Biology, 30(2), pp. 197–206. doi:Doi 10.1016/S0163-1047(80)91077-8

Pantazis, C. B., James, M. H., Bentzley, B. S., & Aston-Jones, G. (2019). The number of lateral hypothalamus orexin/hypocretin neurons contributes to individual differences in cocaine demand. Addict Biol, p e12795. doi:10.1111/adb.12795

Parise, E. M., Lilly, N., Kay, K., Dossat, A. M., Seth, R., Overton, J. M., & Williams, D. L. (2011). Evidence for the role of hindbrain orexin-1 receptors in the control of meal size. Am J Physiol Regul Integr Comp Physiol, 301(6), pp. R1692–1699. doi:10.1152/ajpregu.00044.2011

Paul, M. J., Terranova, J. I., Probst, C. K., Murray, E. K., Ismail, N. I., & de Vries, G. J. (2014). Sexually dimorphic role for vasopressin in the development of social play. Frontiers in Behavioral Neuroscience, 8. doi:10.3389/Fnbeh.2014.00058

Pellis, S. M., & Iwaniuk, A. N. (2000). Comparative analyses of the role of postnatal development on the expression of play fighting. Dev Psychobiol, 36(2), pp. 136–147.

Pellis, S. M., & Mckenna, M. M. (1992). Intrinsic and xxtrinsic influences on play fighting in rats - Effects of dominance, partners playfulness, temperament and neonatal exposure to testosterone propionate. Behavioural Brain Research, 50(1-2), pp. 135–145. doi:Doi 10.1016/S0166-4328(05)80295-5

Perez-Leighton, C. E., Boland, K., Billington, C. J., & Kotz, C. M. (2013). High and low activity rats: elevated intrinsic physical activity drives resistance to diet-induced obesity in non-bred rats. Obesity (Silver Spring), 21(2), pp. 353–360. doi:10.1002/oby.20045

Petrovich, G. D., Hobin, M. P., & Reppucci, C. J. (2012). Selective Fos induction in hypothalamic orexin/hypocretin, but not melanin-concentrating hormone neurons, by a learned food-cue that stimulates feeding in sated rats. Neuroscience, 224, pp. 70–80. doi:10.1016/j.neuroscience.2012.08.036

Peyron, C., Tighe, D. K., van den Pol, A. N., de Lecea, L., Heller, H. C., Sutcliffe, J. G., & Kilduff, T. S. (1998). Neurons containing hypocretin (orexin) project to multiple neuronal systems. J Neurosci, 18, pp. 9996–10015.

Poole, T. B., & Fish, J. (1976). An investigation of individual, age and sexual differences in the play of Rattus norvegicus (Mammalia: Rodentia). Journal of Zoology, 172(2), pp. 249–259.

Reppucci, C. J., Gergely, C. K., & Veenema, A. H. (2018). Activation patterns of vasopressinergic and oxytocinergic brain regions following social play exposure in juvenile male and female rats. J Neuroendocrinol doi:10.1111/jne.12582

Richardson, K. A., & Aston-Jones, G. (2012). Lateral hypothalamic orexin/hypocretin neurons that project to ventral tegmental area are differentially activated with morphine preference. J Neurosci, 32(11), pp. 3809–3817. doi:10.1523/JNEUROSCI.3917-11.2012

Sakurai, T., Amemiya, A., Ishii, M., Matsuzaki, I., Chemelli, R. M., Tanaka, H., … Yanagisawa, M. (1998). Orexins and orexin receptors: a family of hypothalamic neuropeptides and G protein-coupled receptors that regulate feeding behavior. Cell, 92(4), pp. 573–585. doi:10.1016/s0092-8674(00)80949-6

Saper, C. B. (2006). Staying awake for dinner: hypothalamic integration of sleep, feeding, and circadian rhythms. Prog Brain Res, 153, pp. 243–252. doi: 10.1016/S0079-6123(06)53014-6

Sawai, N., Ueta, Y., Nakazato, M., & Ozawa, H. (2010). Developmental and aging change of orexin-A and-B immunoreactive neurons in the male rat hypothalamus. Neuroscience letters, 468(1), pp. 51–55. doi: 10.1016/j.neulet.2009.10.061

Schmitt, O., Usunoff, K. G., Lazarov, N. E., Itzev, D. E., Eipert, P., Rolfs, A., & Wree, A. (2012). Orexinergic innervation of the extended amygdala and basal ganglia in the rat. Brain Struct Funct, 217(2), pp. 233–256. doi:10.1007/s00429-011-0343-8

Sharko, A. C., Fadel, J. R., Kaigler, K. F., & Wilson, M. A. (2017). Activation of orexin/hypocretin neurons is associated with individual differences in cued fear extinction. Physiol Behav, 178, pp. 93–102. doi:10.1016/j.physbeh.2016.10.008

Siegel, P. S. (1961). Food intake in rat in relation to dark-light cycle. Journal of comparative and physiological Psychology, 54(3), pp. 294–301. doi:10.1037/h0044787

Silva, E. S. D., Flores, R. A., Ribas, A. S., Taschetto, A. P., Faria, M. S., Lima, L. B., … Paschoalini, M. A. (2017). Injections of the of the alpha1-adrenoceptor antagonist prazosin into the median raphe nucleus increase food intake and Fos expression in orexin neurons of free-feeding rats. Behav Brain Res, 324, pp. 87–95. doi:10.1016/j.bbr.2017.02.021

Siviy, S. M. (2016). A brain motivated to play: insights into the neurobiology of playfulness. Behavior, 153(6-7), pp. 819–844. doi:10.1163/1568539X-00003349

Siviy, S. M., Crawford, C. A., Akopian, G., & Walsh, J. P. (2011). Dysfunctional play and dopamine physiology in the Fischer 344 rat. Behavioural Brain Research, 220(2), pp. 294–304. doi:10.1016/j.bbr.2011.02.009

Siviy, S. M., Love, N. J., DeCicco, B. M., Giordano, S. B., & Seifert, T. L. (2003). The relative playfulness of juvenile Lewis and Fischer-344 rats. Physiology & Behavior, 80(2-3), pp. 385–394. doi:10.1016/j.physbeh.2003.09.002

Spinka, M., Newberry, R. C., & Bekoff, M. (2001). Mammalian play: Training for the unexpected. Quarterly Review of Biology, 76(2), pp. 141–168. doi:10.1086/393866

Stephan, F. K., & Zucker, I. (1972). Circadian rhythms in drinking behavior and locomotor activity of rats are eliminated by hypothalamic lesions. Proc Natl Acad Sci U S A, 69(6), pp. 1583–1586. doi:10.1073/pnas.69.6.1583

Swanson, L. W. (2018). Brain maps 4.0-Structure of the rat brain: An open access atlas with global nervous system nomenclature ontology and flatmaps. J Comp Neurol, 526(6), pp. 935–943. doi:10.1002/cne.24381

Swanson, L. W., Sanchez-Watts, G., & Watts, A. G. (2005). Comparison of melanin-concentrating hormone and hypocretin/orexin mRNA expression patterns in a new parceling scheme of the lateral hypothalamic zone. Neurosci Lett, 387(2), pp. 80–84. doi:10.1016/j.neulet.2005.06.066

Taheri, S., Mahmoodi, M., Opacka-Juffry, J., Ghatei, M. A., & Bloom, S. R. (1999). Distribution and quantification of immunoreactive orexin A in rat tissues. FEBS letters, 457(1), pp. 157–161. doi:10.1016/s0014-5793(99)01030-3

Taylor, G. T. (1980). Fighting in Juvenile Rats and the Ontogeny of Agonistic Behavior. Journal of comparative and physiological Psychology, 94(5), pp. 953–961. doi:10.1037/H0077816

Trivedi, P., Yu, H., MacNeil, D. J., Van der Ploeg, L. H., & Guan, X. M. (1998). Distribution of orexin receptor mRNA in the rat brain. FEBS Lett, 438(1-2), pp. 71–75. doi:10.1016/s0014-5793(98)01266-6

Tsujino, N., & Sakurai, T. (2009). Orexin/hypocretin: a neuropeptide at the interface of sleep, energy homeostasis, and reward system. Pharmacol Rev, 61(2), pp. 162–176. doi:10.1124/pr.109.001321

van den Berg, C. L., Hol, T., Van Ree, J. M., Spruijt, B. M., Everts, H., & Koolhaas, J. M. (1999). Play is indispensable for an adequate development of coping with social challenges in the rat. Developmental Psychobiology, 34(2), pp. 129–138. doi:10.1002/(Sici)1098-2302(199903)34:2<129::Aid-Dev6>3.0.Co;2-L

Vanderschuren, L. J., Achterberg, E. J., & Trezza, V. (2016). The neurobiology of social play and its rewarding value in rats. Neurosci Biobehav Rev, 70, pp. 86–105. doi:10.1016/j.neubiorev.2016.07.025

Vanderschuren, L. J. M. J., Niesink, R. J. M., & VanRee, J. M. (1997). The neurobiology of social play behavior in rats. Neuroscience and Biobehavioral Reviews, 21(3), pp. 309–326. doi:10.1016/S0149-7634(96)00020-6

Veenema, A. H., Bredewold, R., & de Vries, G. J. (2013). Sex-specific modulation of juvenile social play by vasopressin. Psychoneuroendocrinology, 38(11), pp. 2554–2561. doi:10.1016/j.psyneuen.2013.06.002

Vittoz, N. M., Schmeichel, B., & Berridge, C. W. (2008). Hypocretin /orexin preferentially activates caudomedial ventral tegmental area dopamine neurons. Eur J Neurosci, 28(8), pp. 1629–1640. doi:10.1111/j.1460-9568.2008.06453.x

Willie, J. T., Chemelli, R. M., Sinton, C. M., Tokita, S., Williams, S. C., Kisanuki, Y. Y., … Yanagisawa, M. (2003). Distinct narcolepsy syndromes in Orexin receptor-2 and Orexin null mice: molecular genetic dissection of Non-REM and REM sleep regulatory processes. Neuron, 38(5), pp. 715–730. doi: 10.1016/s0896-6273(03)00330-1

Wu, M. F., Nienhuis, R., Maidment, N., Lam, H. A., & Siegel, J. M. (2011). Cerebrospinal fluid hypocretin (orexin) levels are elevated by play but are not raised by exercise and its associated heart rate, blood pressure, respiration or body temperature changes. Arch Ital Biol, 149(4), pp. 492–498. doi:10.4449/aib.v149i4.1315

Yamamoto, Y., Ueta, Y., Hara, Y., Serino, R., Nomura, M., Shibuya, I., … & Yamashita, H. (2000). Postnatal development of orexin/hypocretin in rats. Molecular brain research, 78(1-2), pp. 108–119. doi:10.1016/s0169-328x(00)00080-2

Zheng, H., Patterson, L. M., & Berthoud, H. R. (2007). Orexin signaling in the ventral tegmental area is required for high-fat appetite induced by opioid stimulation of the nucleus accumbens. J Neurosci, 27(41), pp. 11075–11082. doi:10.1523/JNEUROSCI.3542-07.2007

